# Identifying differences in bile acid pathways for cholesterol clearance in Alzheimer’s disease using metabolic networks of human brain regions

**DOI:** 10.1101/782987

**Authors:** Priyanka Baloni, Cory C. Funk, Jingwen Yan, James T. Yurkovich, Alexandra Kueider-Paisley, Kwangsik Nho, Almut Heinken, Wei Jia, Siamak Mahmoudiandehkordi, Gregory Louie, Andrew J. Saykin, Matthias Arnold, Gabi Kastenmüller, William J Griffiths, Ines Thiele, AMP-AD consortium, The Alzheimer’s Disease Metabolomics Consortium, Rima Kaddurah-Daouk, Nathan D. Price

## Abstract

Alzheimer’s disease (AD) is the leading cause of dementia, with metabolic dysfunction seen years before the emergence of clinical symptoms. Increasing evidence suggests a role for primary and secondary bile acids, the end-product of cholesterol metabolism, influencing pathophysiology in AD. In this study, we analyzed transcriptomes from 2114 post-mortem brain samples from three independent cohorts and identified that the genes involved in alternative bile acid synthesis pathway was expressed in brain compared to the classical pathway. These results were supported by targeted metabolomic analysis of primary and secondary bile acids measured from post-mortem brain samples of 111 individuals. We reconstructed brain region-specific metabolic networks using data from three independent cohorts to assess the role of bile acid metabolism in AD pathophysiology. Our metabolic network analysis suggested that taurine transport, bile acid synthesis and cholesterol metabolism differed in AD and cognitively normal individuals. Using the brain transcriptional regulatory network, we identified putative transcription factors regulating these metabolic genes and influencing altered metabolism in AD. Intriguingly, we find bile acids from the brain metabolomics whose synthesis cannot be explained by enzymes we find in the brain, suggesting they may originate from an external source such as the gut microbiome. These findings motivate further research into bile acid metabolism and transport in AD to elucidate their possible connection to cognitive decline.

## Introduction

Alzheimer’s disease (AD), the leading cause of dementia, is a progressive, multifactorial disease^1,2^ where the onset and progression of symptoms varies significantly among individuals. Recent studies have shown that metabolic dysfunction is one of the factors associated with neurodegenerative disorders^3,4^. Various physiological processes such as lipid metabolism, immune function, amyloid precursor protein metabolism, oxidative stress, neurotransmitter function as well as mitochondrial functions are altered in AD that can affect metabolism^5,6,7^. Interest in the transport of biochemical compounds between the brain and the gut and their possible role in regulating metabolic changes centrally and peripherally has increased recently across several neurodegenerative diseases^8,9^. There is increasing evidence to suggest a role in AD for primary and secondary bile acids^7,10,11^. Bile acids are amphipathic molecules and primary bile acids are derived from cholesterol mostly in the liver, whereas secondary bile acids are typically produced by bacteria in the gut^12^. Increased levels of secondary bile acids and ratios to their primary bile acid educts have been linked to AD and cognitive decline^7^.

Cholesterol metabolism and transport have been studied extensively and are clearly linked with AD^13,1,14^. Cholesterol clearance leads to production of bile acids that carry out lipid absorption, cholesterol homeostasis and also function as signaling molecules^15^. Primary bile acids such as cholic acid and chenodeoxycholic acid are synthesized as a result of cholesterol efflux and then conjugated with glycine or taurine for secretion into bile and later metabolized by gut bacteria^12^. There are two major bile acid biosynthetic pathways: the classical pathway (neutral pathway) and the alternative pathway (acidic pathway). The classical pathway in mammalian liver is initiated by cholesterol 7α-hydroxylase (*CYP7A1)* and subsequently requires 12α-hydroxylase (*CYP8B1*) amongst numerous other enzymes for synthesis of cholic acid, whereas chenodeoxycholic acid is produced in the absence of *CYP8B1*^*16*^. Sterol 27-hydroxylase (*CYP27A1*) is required for the initiation of alternative bile acid pathway^17^. In the brain, sterol 24-hydroxylase (*CYP46A1*) converts cholesterol to 24S-hydroxycholesterol (systematic name cholest-5-en-3β,24S-diol), and subsequent 7α-hydroxylation is carried out by 24-hydroxycholesterol 7α-hydroxylase (*CYP39A1*)^18^ (Figure 1). Studies in human and mouse brain samples, as well as cell lines have shown that bile acids can cross the blood-brain barrier and bind to nuclear receptors, causing physiological changes^19,20^. There is limited information on the role of bile acids in human brain and their association with cognitive decline in AD pathophysiology. Systematic analysis of omics data derived from blood and post-mortem brain samples of AD and cognitively normal (CN) or control individuals has the potential to identify differences in cholesterol and bile acid metabolism and how they contribute to AD pathogenesis.

**Figure 1:**
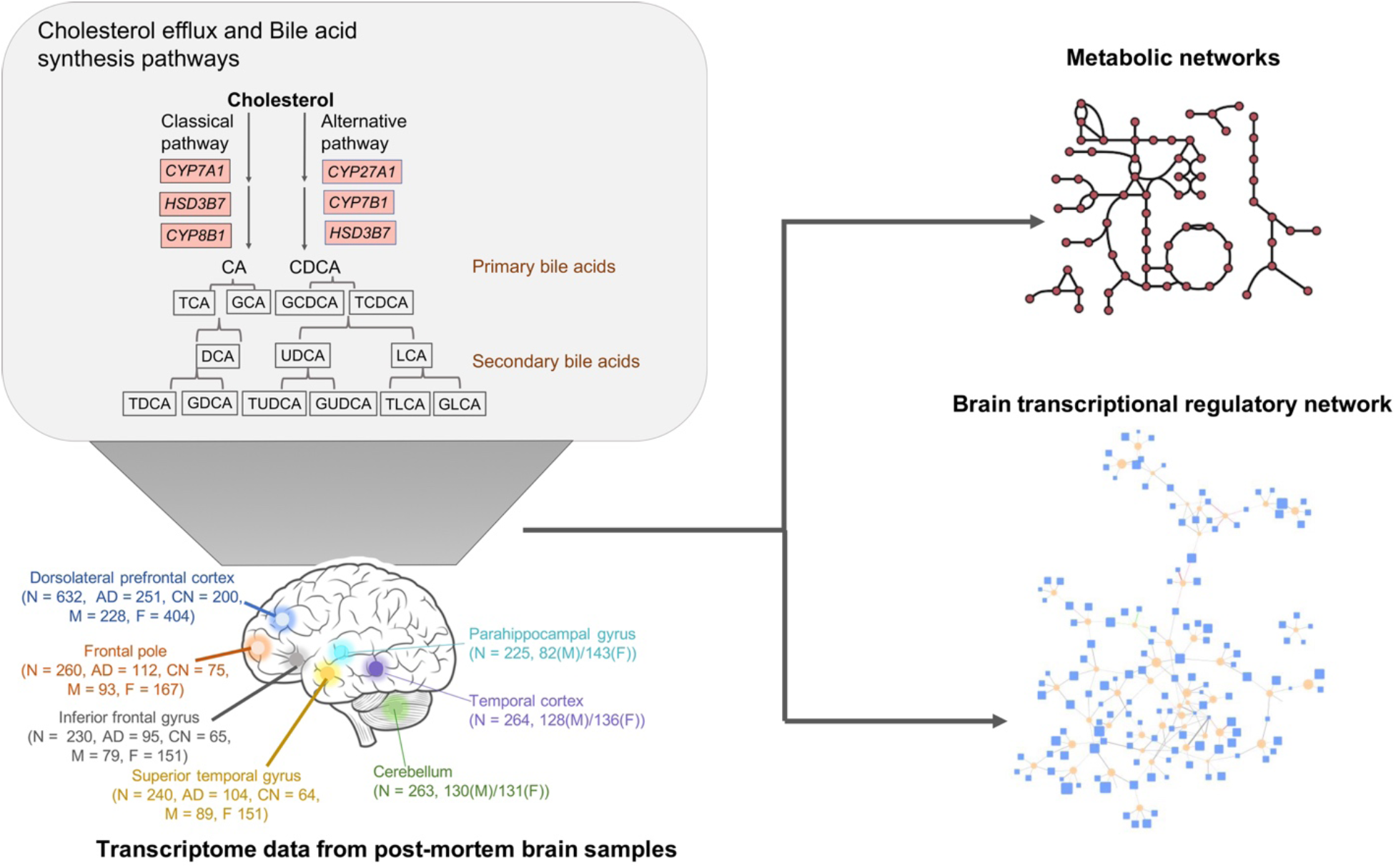
Graphical overview of analyses described herein to study altered cholesterol and bile acid metabolism in AD. Numbers of samples from each brain region are indicated along with AD and control samples and Male/Female breakdown in parentheses. We analyzed the metabolic networks for each sample and also used regulatory models to identify important transcription factors regulating cholesterol and bile acid metabolism.

In this study, we analyzed large number of transcriptome data from the Religious Orders Study and Memory and Aging Project (ROSMAP), Mayo Clinic and the Mount Sinai Brain Bank that had a total of 2114 post-mortem brain samples from seven different brain regions. We reconstructed metabolic networks using the data from these three cohorts and studied the role of circulating bile acids that may contribute to AD and altered cholesterol metabolism in these individuals. We also generated targeted metabolomics data of primary and secondary bile acids from post-mortem brain samples of 111 AD patients and controls.

Various genomic studies have reported transcriptional regulatory changes in neurodegenerative diseases^21,22^. The biological significance of these transcription factors (TFs) regulating metabolic changes is not completely understood. The brain-specific metabolic and transcriptional regulatory networks proved useful in identifying candidate metabolites and genes involved in the disease manifestation. A schematic representation of the study is represented in Figure 1. Our study used the following approaches to investigate the role of bile acids in AD:

i. Transcriptional profiling of genes from post-mortem brain samples that are involved in cholesterol and bile acid metabolism.
ii. Reconstruction and analysis of genome-scale metabolic networks of various brain regions to identify genes and reactions that are significant in AD vs CN.
iii. Transcriptional regulatory network analysis of brain samples to predict candidate TFs regulating metabolically important genes.

In summary, our study addresses an important need to better understand potential roles for bile acids in AD pathophysiology.

## Results

In recent studies, cytotoxic and neuroprotective bile acids were identified in AD and their probable link to cognitive decline in the individuals was reported^7,23^. To further investigate the role of primary and secondary bile acids in AD and CN individuals, we analyzed 2114 post-mortem brain samples from three independent cohorts for seven brain regions (Table 1) and selected genes involved in cholesterol and bile acid metabolism.

**Table 1:** Details of brain region-specific samples from three different cohorts. The total number of post-mortem brain samples for different regions, method of preparation of RNA-sequencing library preparation, number of samples for different pathologies, gender and APOE status of the individuals is described in the table.

| Brain regions | Source | Number of brain samples | RNA-Seq Library prep method | Pathologies | Gender | APOE allele |
| --- | --- | --- | --- | --- | --- | --- |
| Cerebellum (CER) | Mayo Clinic, University of Florida, Institute for Systems Biology | 263 | poly-A enriched | AD (79), control (72), NA (2), Pathologic aging (28), PSP (82) | Male (130), Female (131), NA (2) | $\epsilon 2/\epsilon 2$ (1), $\epsilon 2/\epsilon 3$ (29), $\epsilon 2/\epsilon 4$ (2), $\epsilon 3/\epsilon 3$ (159), $\epsilon 3/\epsilon 4$ (64), $\epsilon 4/\epsilon 4$ (6), NA (2) |
| Temporal cortex (TCX) | Mayo Clinic, University of Florida, Institute for Systems Biology | 264 | poly-A enriched | AD (80), control (73), Pathologic aging (30), PSP (81) | Male (128), Female (136) | $\epsilon 2/\epsilon 2$ (1), $\epsilon 2/\epsilon 3$ (33), $\epsilon 2/\epsilon 4$ (2), $\epsilon 3/\epsilon 3$ (157), $\epsilon 3/\epsilon 4$ (62), $\epsilon 4/\epsilon 4$ (9) |
| Prefrontal cortex (FC) | Religions Orders Study and Memory and Aging Project (ROSMAP) | 632 | strand-specific | AD (251), MCI (169), NCI (200), Other dementia (12) | Male (228), Female (404) | $\epsilon 2/\epsilon 2$ (5), $\epsilon 2/\epsilon 3$ (83), $\epsilon 2/\epsilon 4$ (17), $\epsilon 3/\epsilon 3$ (382), $\epsilon 3/\epsilon 4$ (139), $\epsilon 4/\epsilon 4$ (6), NA (1) |
| Frontal Pole (FP) | Mount Sinai Brain Bank (MSBB) | 260 | Ribo-zero | AD (112), Normal (75), Possible D (38), Probable AD (35) | Male (93), Female (167) | $\epsilon 2/\epsilon 2$ (2), $\epsilon 2/\epsilon 3$ (18), $\epsilon 2/\epsilon 4$ (1), $\epsilon 3/\epsilon 3$ (88), $\epsilon 3/\epsilon 4$ (49), $\epsilon 4/\epsilon 4$ (3), NA (99) |
| Superior temporal gyrus (STG) | Mount Sinai Brain Bank (MSBB) | 240 | Ribo-zero | AD (104), Normal (64), Possible AD (36), Probable AD (36) | Male (89), Female (151) | $\epsilon 2/\epsilon 2$ (1), $\epsilon 2/\epsilon 3$ (17), $\epsilon 2/\epsilon 4$ (1), $\epsilon 3/\epsilon 3$ (81), $\epsilon 3/\epsilon 4$ (45), $\epsilon 4/\epsilon 4$ (3), NA (92) |
| Inferior frontal gyrus (IFG) | Mount Sinai Brain Bank (MSBB) | 230 | Ribo-zero | AD (95), Normal (65), Possible AD (36), Probable AD (34) | Male (79), Female (151) | $\epsilon 2/\epsilon 2$ (2), $\epsilon 2/\epsilon 3$ (15), $\epsilon 2/\epsilon 4$ (1), $\epsilon 3/\epsilon 3$ (77), $\epsilon 3/\epsilon 4$ (41), $\epsilon 4/\epsilon 4$ (3), NA (91) |
| Parahippocamal gyrus (PHG) | Mount Sinai Brain Bank (MSBB) | 225 | Ribo-zero | AD (97), Normal (68), Possible AD (28), Probable AD (32) | Male (82), Female (143) | $\epsilon 2/\epsilon 2$ (2), $\epsilon 2/\epsilon 3$ (16), $\epsilon 2/\epsilon 4$ (1), $\epsilon 3/\epsilon 3$ (71), $\epsilon 3/\epsilon 4$ (36), $\epsilon 4/\epsilon 4$ (3), NA (96) |

### Transcriptomic analysis of enzyme-encoding genes associated with bile acid metabolism

Bile acids are products of cholesterol metabolism. To identify cholesterol and bile acid genes that are expressed in brain, we curated a list of regulators, transporters and biosynthesis genes in these three independent cohorts. Cholesterol biosynthesis regulators *SREBF1* and *SREBF2* were expressed in post-mortem brain samples and recent studies have identified variants of SREBP2, the protein encoded by *SREBF2*, and their probable link with AD^14,24,25^. Expression of genes involved in cholesterol transport *ABCA1, ABCA5, ABCA7, APOE, LPL* and *LCAT* and members of the LDLR gene family (*LDLR, VLDLR, LRP1, LRP2, LRP4, LRP5, LRP6, LRP8, LRAD3*) in the brain samples suggests active transport of cholesterol and cholesterol homeostasis in brain (Figure 2). *ABCA7*, a cholesterol transporter, belonging to the class of ATP-binding cassette transporters that has been identified as a risk factor for late-onset of AD^17^, is not found in the existing KEGG pathways and was manually curated into our models. *ABCA7* was found to be expressed in the post-mortem brain samples. We also probed into genes encoding for receptors linked with the classical and alternative bile acid pathway and found expression of *PPARA, PPARG, LXR*α*/β, RAR* and RXRs (*RXRA, RXRB, RXRG*) in the samples but no evidence of expression of FXR.

**Figure 2:**
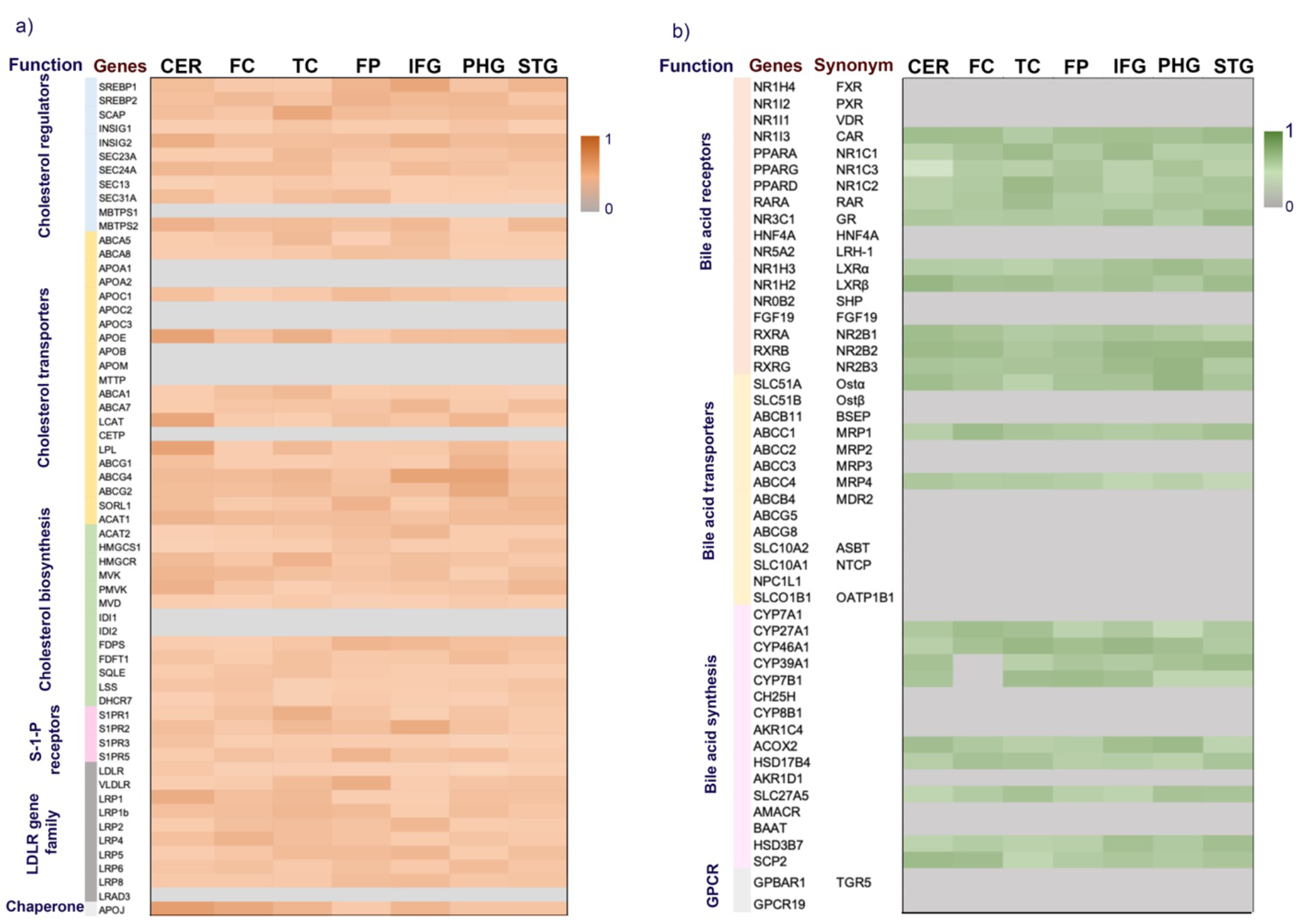
Heatmap for genes involved in (a) cholesterol and (b) bile acid metabolism. The color gradient is based on ubiquity score calculated for the genes and gray color represents genes having no expression data on the brain regions from three cohorts. Brain regions represented in the plot are cerebellum (CER), prefrontal cortex (FC), temporal cortex (TC), frontal pole (FP), inferior frontal gyrus (IFG), parahippocampal gyrus (PHG) and superior temporal gyrus (STG). The function of genes is indicated on the left side of each heatmap.

We observed consistent expression of *CYP27A1* and *CYP7B1*, which are involved in the initial steps of the alternative bile acid pathway depicted in Figure 3, from the analysis of transcriptomic data of post-mortem brain samples from three independent cohorts (Supplementary file 1). In the figure, the bile acids have been marked as cytotoxic and neuroprotective^7,23^, but all bile acids become toxic at elevated concentrations because of their ability to solubilize membranes^23^. We did not observe expression of *CYP7A1* and *CYP8B1*, suggesting that the classical bile acid biosynthesis pathway is not prevalent in the brain samples. The classical pathway is known to be most active in the liver^26^. It has been reported that neural cholesterol clearance through bile acid synthesis is mediated by *CYP46A1* and subsequently by *CYP39A1* in the liver, leading to synthesis of chenodeoxycholic acid^27^. In addition to genes involved in the alternative bile acid pathway, we also observed expression of brain-specific *CYP46A1* and *CYP39A1* genes in all the cohorts. This analysis suggested that the brain utilizes an alternative and neural cholesterol clearance pathway of bile acid synthesis^27,11^ and not the classical pathway^23^.

**Figure 3:**
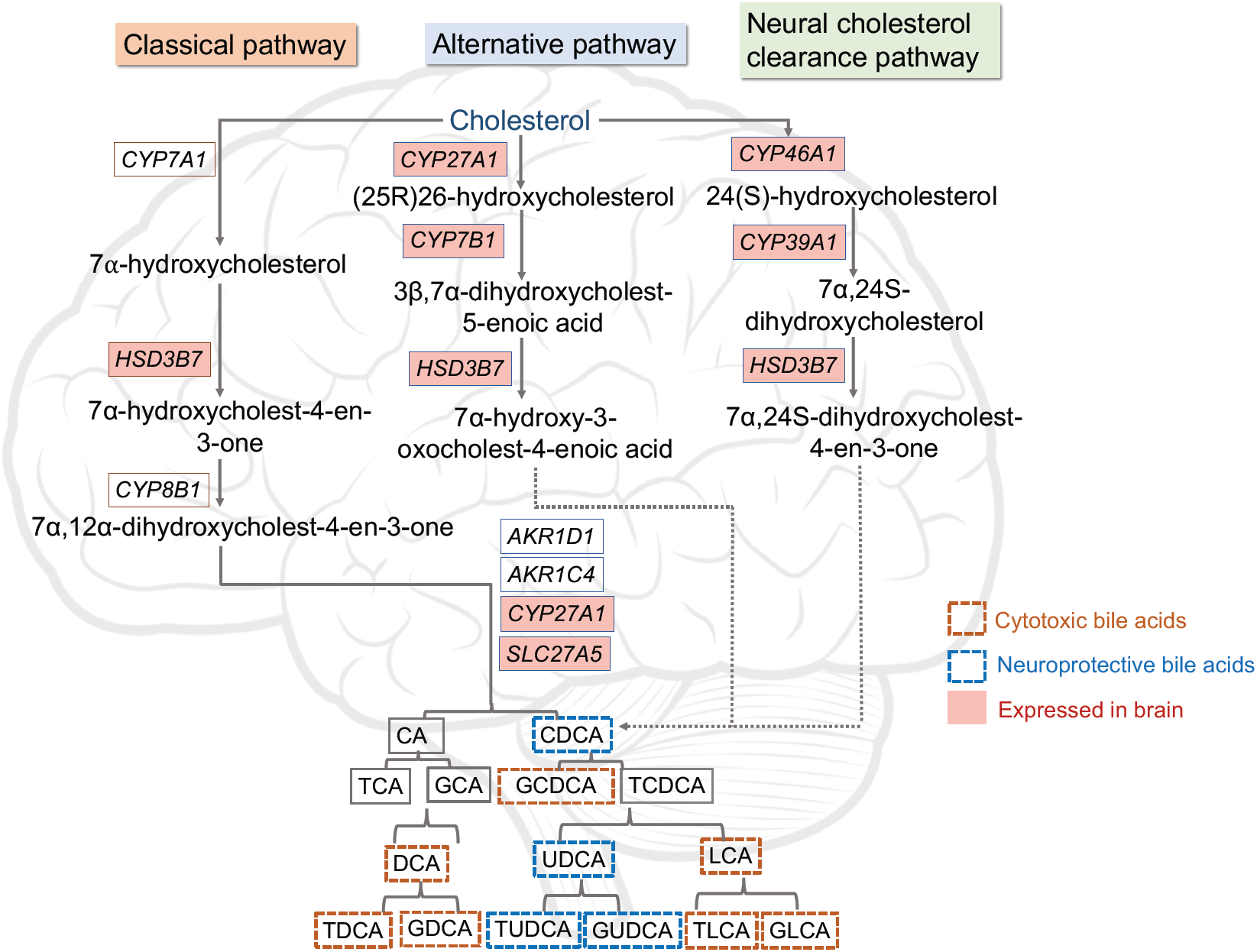
Schematic representation of bile acid synthesis pathway in humans. The order of enzymatic reactions can vary. Genes expressed in brain samples from our analysis are highlighted in pink. Based on the results from^7^, bile acids have been marked as neuroprotective or cytotoxic.

### Metabolomics analysis of post-mortem brain samples to identify levels of primary and secondary bile acids

Bile acids were quantified from 111 post-mortem brain samples from the dorsolateral prefrontal cortex of AD, MCI and CN individuals in the ROSMAP study (https://www.synapse.org/#!Synapse:syn10235594) (Supplementary file 2). Although the genes involved in production of cholic acid were not expressed in the brain samples, detection of cholic acid from the metabolomics analysis suggested that cholic acid might enter the brain from the periphery as previously shown in other studies^20,28,29^. We compared the levels of primary and secondary bile acids in individuals with CERAD score of 1-4, where 1, 2, 3 and 4 indicate definitive AD, probable AD, possible AD and no evidence of AD, respectively. The ratio of primary conjugated and secondary bile acids with respect to cholic acid (CA) showed that deoxycholic acid (DCA), lithocholic acid (LCA), glycochenodeoxycholate (GCDCA), chenodeoxycholic acid (CDCA), taurodeoxycholic acid (TDCA), glycodeoxycholic acid (GDCA), ursodeoxycholic acid (UDCA), allolithocholate (alloLCA) and taurocholic acid (TCA) were higher in individuals with AD (CERAD score 1-3) compared to controls (Figure 4). Similar results were reported in the serum metabolomics samples of AD and CN individuals^7,10^. Allo-cholic acid (ACA) is a steroid bile acid has been studied in the context of signaling mechanisms related to differentiation, proliferation or apoptosis of hepatocytes^30^. The CDCA:CA ratio was calculated and it showed the higher value for AD compared to CN individuals in the study (Supplementary file 3). This finding suggests that the alternative bile acid pathway is more active in AD versus CN individuals. Also, the higher ratio of primary bile acid like TCA and secondary bile acids such as DCA, LCA, TDCA and GDCA in AD individuals indicated that these bile acids may be associated with cognitive function.

**Figure 4:**
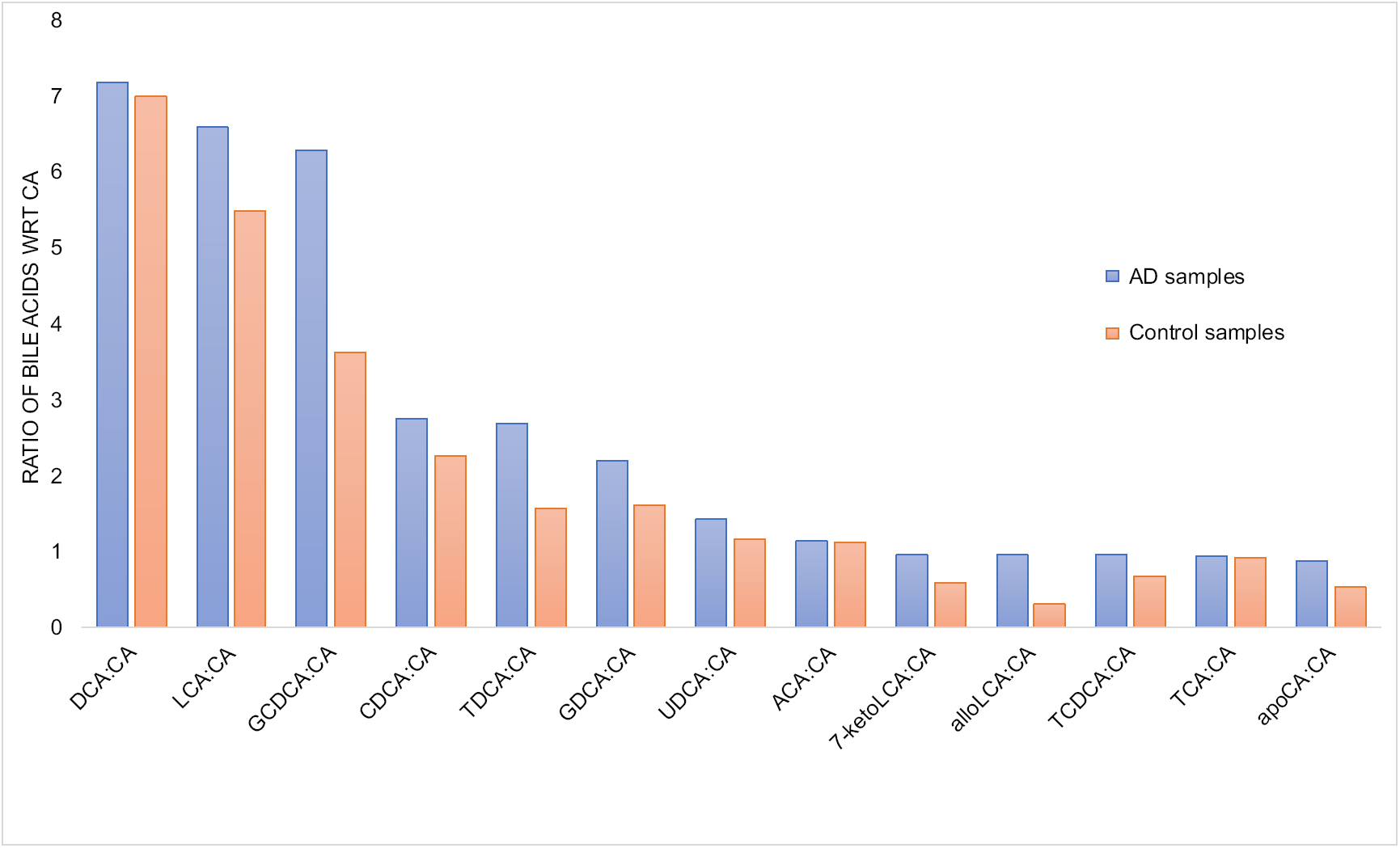
Bar plots representing ratio of bile acids with respect to cholic acid (primary bile acid) measured from 111 brain samples from ROSMAP study. Blue bars represent AD samples and light orange bars represent control samples.

The primary BAs are conjugated with glycine or taurine for secretion into bile^17^. In addition to the primary and secondary BAs, we also measured levels of taurine in serum samples in AD and CN individuals. In the serum, we observed that AD patients had higher levels of serum taurine compared to controls. Taurine is required for conjugation of primary and secondary bile acids. This is an interesting observation and we need to explore the transport and physiological levels of taurine in the brains of individuals with AD.

### Metabolic reconstruction of brain regions and pathway-level analysis

We reconstructed metabolic networks for brain region-specific samples in the three independent cohorts. The seven brain regions in this study included cerebellum (CER), prefrontal cortex (FC), temporal cortex (TC), frontal pore (FP), inferior frontal gyrus (IFG), parahippocampal gyrus (PHG) and superior temporal gyrus (STG). We used transcriptome data from post-mortem brain samples for reconstructing metabolic networks (see Methods for more details). The brain region-specific metabolic networks consisted of ∼5600-6300 reactions, 2800-4000 metabolites and each model had genes varying from 1500-1757 in these networks. Supplementary Figure 1a provides information of the numbers of reactions, metabolites and genes present in each of the brain region-specific networks and Supplementary Figure 1b compares the gene content overlaps across each of these networks. We have made the detailed content of all of these models available to the scientific community (Supplementary file 4-10).

We tested each model using 17 brain-specific in silico tests meant to mimic experimental evidence of metabolic functions in the brain (’metabolic tasks’) (Supplementary file 11), that were obtained from a recently published work on human reconstruction^31^. These metabolic tasks represent a set of reactions that are brain-specific, and the metabolic networks generated passed 50-70% of the tasks (see Methods section). The metabolic tasks are listed in Supplementary file 11 and the models have been provided in SBML in Supplementary file 12-18. In addition to generating brain region-specific metabolic networks, we also used the transcriptome data of 2114 post-mortem brain samples and obtained personalized networks for each sample in the study. Out of 2114 brain samples, 818 samples corresponded to individuals with AD, 138 possible AD, 137 probable AD and 617 controls. The dataset also consisted of 12 samples from other dementias, 163 samples with progressive supranuclear palsy, 58 samples with pathologic aging and 2 samples that were uncharacterized. From our metabolic networks, we identified 518 reactions that were involved in cholesterol metabolism, bile acid synthesis and transport of bile acids between different compartments in the metabolic networks. The personalized metabolic networks had distinct set of bile acid reactions active in the brain regions (details in the Methods section). Since the post-mortem brain samples for the brain regions were collected by three independent cohorts having different sequencing protocols and depth, the flux results were analyzed separately for these cohorts. The data suggest that the cerebellum and temporal cortex have similar sets of bile acid reactions that can be active in the personalized metabolic networks (Figure 5).

**Figure 5:**
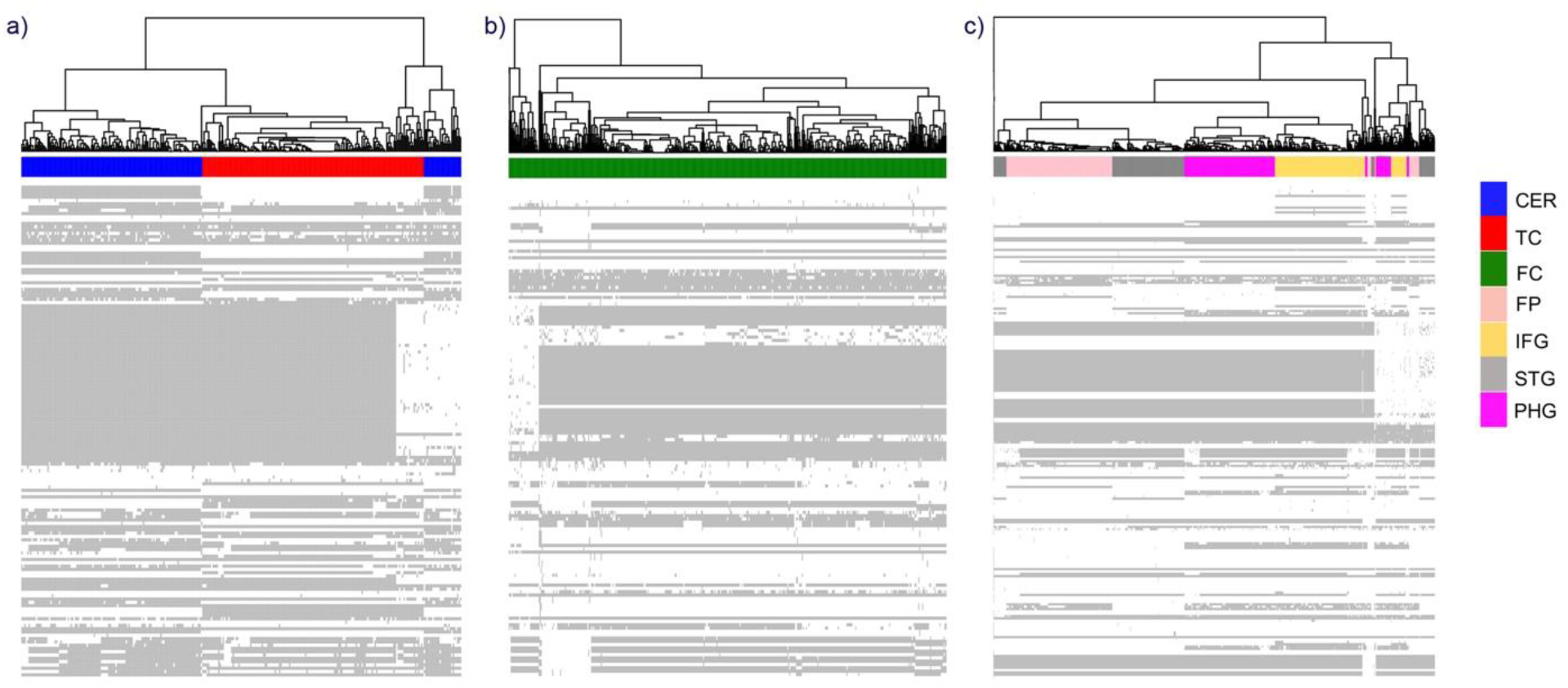
Clustergram for 518 reactions involved in bile acid metabolism in (a) Mayo clinic cohort (cerebellum and temporal cortex), (b) ROSMAP cohort (frontal cortex) and (c) Mount Sinai Brain Bank (frontal pole, inferior temporal gyrus, superior temporal gyrus and parahippocampal gyrus). The rows correspond to bile acid reactions in the network and the columns are colored based on the brain regions.

We analyzed the reaction fluxes and found similar set of bile acid reactions carrying fluxes in metabolic networks of these independent cohorts. We used this information to carry out statistical analysis and identify reactions that are significantly different (p-value < 0.05) in brain regions of AD versus the CN individuals as well as identify reactions that were significant in males versus females with AD. We found reactions carrying out transport of taurine and cholesterol were significant in the dorsolateral prefrontal cortex, temporal cortex and parahippocampal gyrus. Taurine is an abundant amino acid present at roughly 1.2 mM in brain^32^. *SLC6A6* (*TAUT*) and *SLC36A1* (*PAT1*) function as taurine transporters and increased transport of taurine across the blood brain barrier (BBB) has been reported for oxidative stress conditions^33^. We found expression of both these genes in the brain transcriptome dataset, suggesting that these genes are expressed in the brain and involved in transport of taurine. Table 2 provides details for significant bile acid reactions in brain regions identified from our analysis.

**Table 2:**
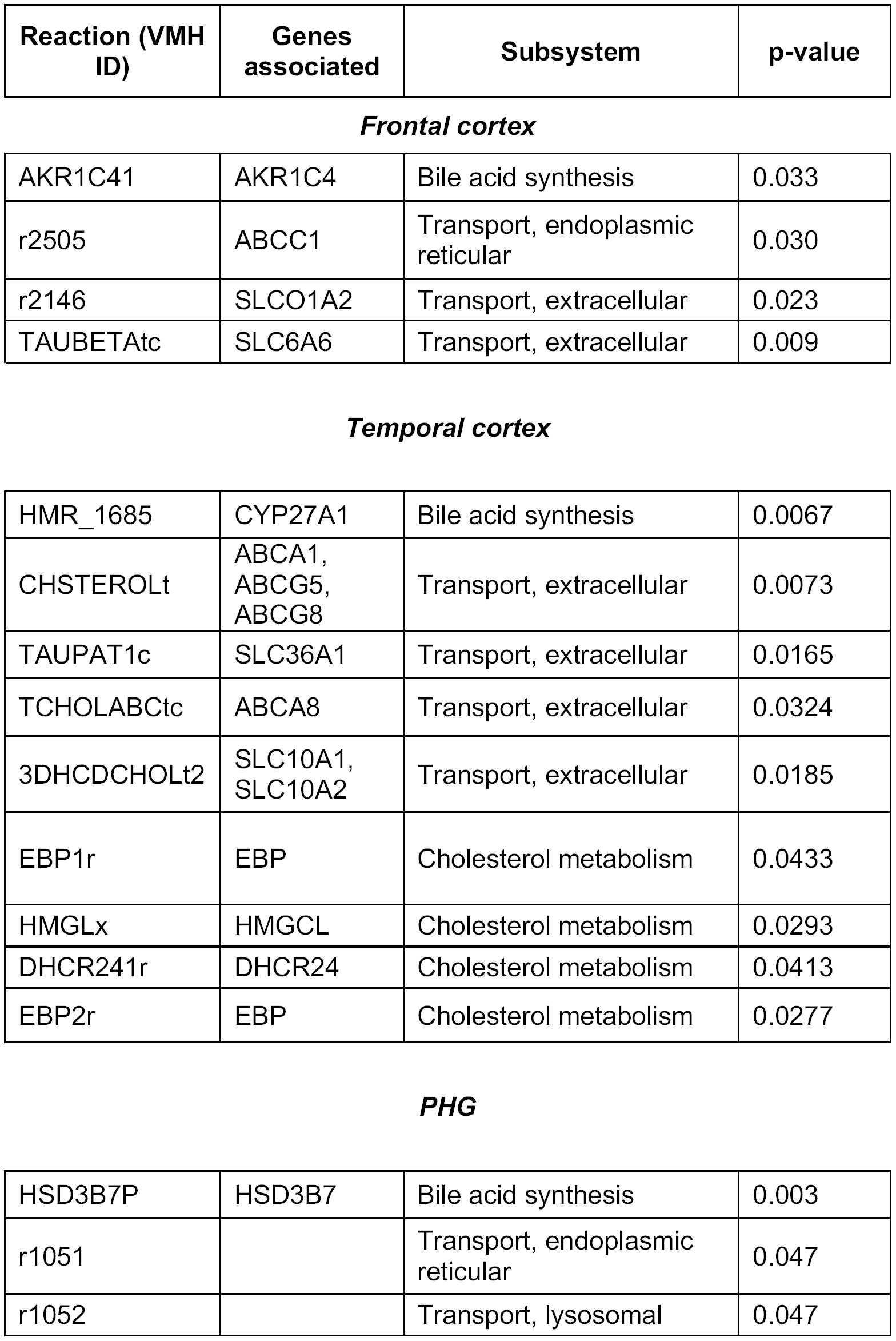
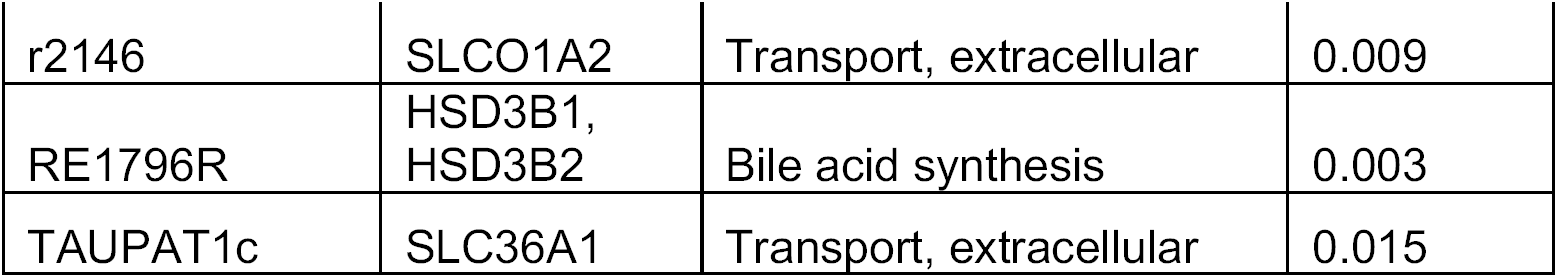
List of bile acid reactions from our metabolic analysis of brain regions. The reactions are represented in their VMH IDs and information related to the genes and subsystems are also shown in the table. P-values calculated by Fisher’s exact test are indicated in the table and only those reactions with p-value < 0.05 are represented here.

| Reaction (VMH ID) | Genes associated | Subsystem | p-value |
| --- | --- | --- | --- |
| **Frontal cortex** |  |  |  |
| AKR1C41 | AKR1C4 | Bile acid synthesis | 0.033 |
| r2505 | ABCC1 | Transport, endoplasmic reticular | 0.030 |
| r2146 | SLCO1A2 | Transport, extracellular | 0.023 |
| TAUBETAtc | SLC6A6 | Transport, extracellular | 0.009 |
| **Temporal cortex** |  |  |  |
| HMR_1685 | CYP27A1 | Bile acid synthesis | 0.0067 |
| CHSTEROLt | ABCA1, ABCG5, ABCG8 | Transport, extracellular | 0.0073 |
| TAUPAT1c | SLC36A1 | Transport, extracellular | 0.0165 |
| TCHOLABCtc | ABCA8 | Transport, extracellular | 0.0324 |
| 3DHCDCHOLt2 | SLC10A1, SLC10A2 | Transport, extracellular | 0.0185 |
| EBP1r | EBP | Cholesterol metabolism | 0.0433 |
| HMGLx | HMGCL | Cholesterol metabolism | 0.0293 |
| DHCR241r | DHCR24 | Cholesterol metabolism | 0.0413 |
| EBP2r | EBP | Cholesterol metabolism | 0.0277 |
| **PHG** |  |  |  |
| HSD3B7P | HSD3B7 | Bile acid synthesis | 0.003 |
| r1051 |  | Transport, endoplasmic reticular | 0.047 |
| r1052 |  | Transport, lysosomal | 0.047 |
| r2146 | SLCO1A2 | Transport, extracellular | 0.009 |
| RE1796R | HSD3B1,<br>HSD3B2 | Bile acid synthesis | 0.003 |
| TAUPAT1c | SLC36A1 | Transport, extracellular | 0.015 |

From our analysis, we identified reactions with *CYP27A1*, required by both the neural cholesterol clearance pathway and the alternative bile acid pathway, as being significant in AD versus CN brains. Other than bile acid synthesis, reactions involving metabolites such as 7α-hydroxycholesterol (Virtual Metabolic Human (VMH, www.vmh.life, REF ID: xol7a), 7α-hydroxy-5β-cholestan-3-one (VMH ID: xol7ah), 3α,7α-dihydroxy-5β-cholestane (VMH ID: xol7ah2) and 7α-hydroxy-cholestene-3-one (VMH ID: xol7aone) were also identified as being significantly different between AD and CN (p-values for these reactions reported in Table 2). Transport of bile acids such as taurolithocholic acid 3-sulfate (VMH ID: HC02198), ursodeoxycholic acid (VMH ID: HC02194), taurocholic acid (VMH ID: tchola) and 3-dehydroxychenodeoxy cholic acid (VMH ID: 3dhchchol) can also be probed further to understand the role of these bile acids in AD. Thus, *in silico* analysis of brain region-specific metabolic models provides insights into reactions that may be involved in metabolic changes in AD that can be validated from experimental data.

### Identifying transcriptional regulators responsible for altered metabolism in AD

Transcription factors are one important aspect of metabolic regulation that operate through adjusting the expression of enzyme-encoding genes. Using a transcriptional regulatory network informed from the same Mayo temporal cortex bulk RNA-seq samples used for the metabolic reconstruction, we identified candidate TFs that interact with metabolic genes in cholesterol and bile acid metabolism. We selectively studied those genes that belonged to reactions that were significantly differentially expressed in AD versus controls, in order to study their role in AD. For example, one gene that came up from our metabolic analysis of AD and controls was emopamil binding protein (*EBP*) (Figure 6). *EBP* is involved in cholesterol metabolism as it is responsible for one of the final steps in the production of cholesterol. Our brain TRN analysis identified *POU6F2, IRF2, SMAD5, GABPA* and *TBR1* as the top candidate TF regulators for EBP. Regulation by these TFs can help in understanding their role in altered cholesterol metabolism in AD, particularly in evaluating the summation of coordinated changes since these TFs of course control other genes as well. *CYP27A1*, as mentioned earlier, is part of the alternative bile acid synthesis pathway and *CREB3L2* and *SOX8* are putative TFs that regulate expression of this gene. *CREB3L2* (cAMP-responsive element binding protein 3-like 2) is induced as a result of ER stress and may function in unfolded protein response signaling in neurons^34^. Other than the metabolism related genes, we also evaluated interactions of bile acid transporters such as *SLC6A6, SLCO1A2, ABCC1, ABCA1, SLC36A1, ABCA8* and their transcriptional regulation. As seen in Figure 6, *SREBF2* was found to interact with *ABCA1* and recently there were reports of variants of SREBP2 that have been linked with AD^25^. Increased *SREBF2* expression leads to higher cholesterol levels and presumably oxysterol and cholestenoic acid levels which are ligands of LXR. The peroxisome proliferator-activated receptors (PPARs) regulate various physiological processes and are expressed in the central nervous system. *PPARA* regulates genes involved in fatty acid metabolism and has been reported to regulate neuronal *ADAM10* expression, in turn affecting proteolysis of amyloid precursor protein^35^. *PPARA* was identified as a putative regulator of *ABCA1* in our brain transcriptional regulatory network. *ABCA1* plays a role in cholesterol metabolism and transport and is a candidate risk gene for late onset Alzheimer’s disease (LOAD)^36^. *SLC6A6*, involved in transport of taurine, was found to be putatively regulated by *STAT1*, a TF reported to play an important role in spatial learning and memory formation^37^, and *RXRG*, that forms heterodimers with retinoic acid (RA), LXRs and vitamin D receptors (VDR) ^38^. Neuronal differentiation 6 (*NEUROD6*) functions in neuronal development, differentiation, and survival in AD^39^. Regulation of *SLC36A1* by *NEUROD6* indicated that this TF plays a role in controlling transport of taurine in the brain. These interactions can be probed further to understand their role in AD pathophysiology.

**Figure 6:**
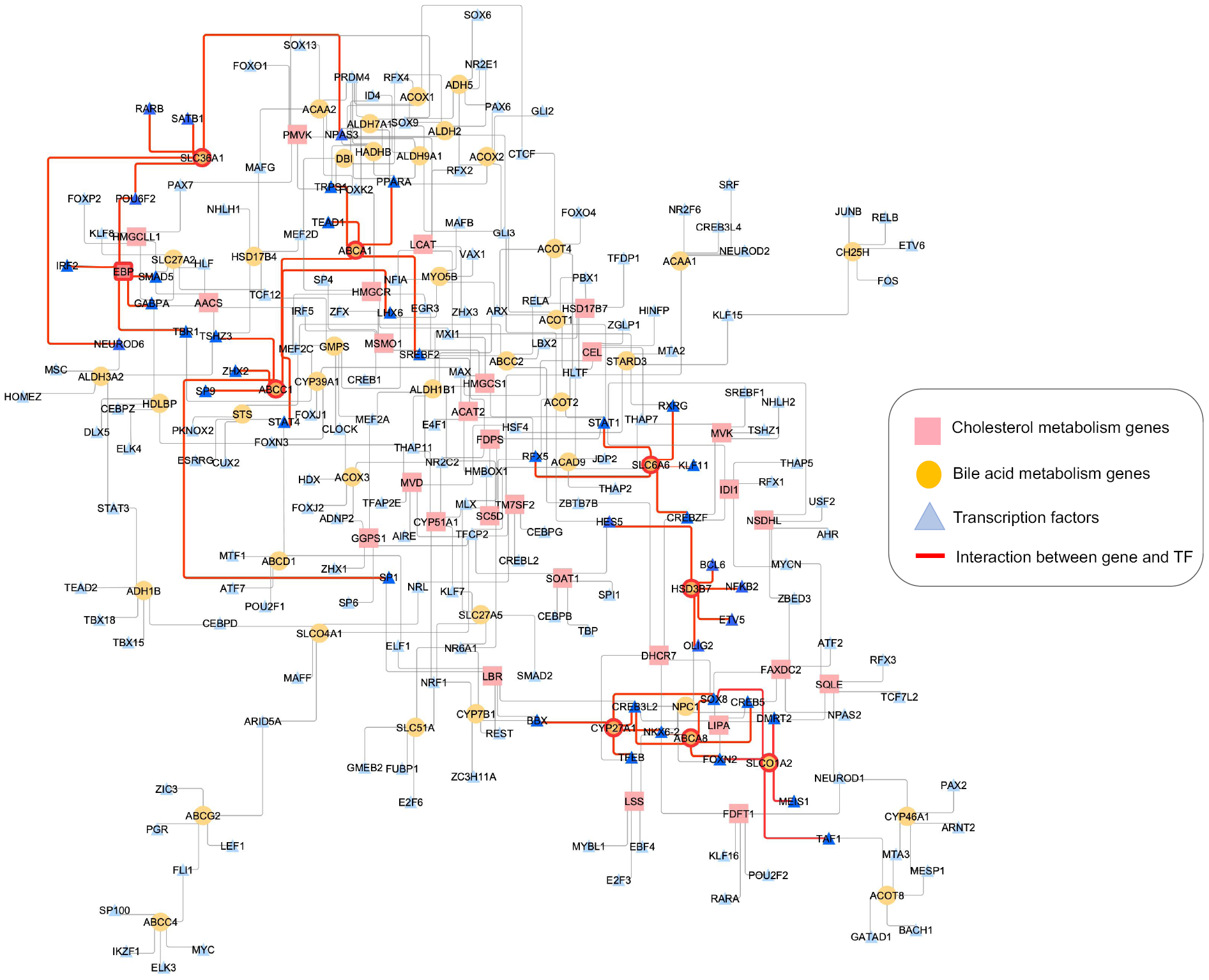
Transcriptional regulatory network of brain highlighting transcription factors and metabolic genes involved in cholesterol and bile acid metabolism. TFs are represented as blue triangles, bile acid metabolism genes as yellow circles and cholesterol metabolism genes as pink rectangles. The significant genes are highlighted with red border and transcription factors in darker shade of blue. Red edges represent interactions between genes and TFs.

In summary, the brain transcriptional regulatory network analysis led to the identification of candidate TFs that regulate genes in cholesterol and BA metabolism, providing clues towards possible roles in bile acid dysregulation in AD.

## Discussion

We carried out a systematic study to identify alterations in cholesterol and bile acid metabolism in AD versus cognitively normal (CN) controls using patient-derived post-mortem transcriptomics and metabolomics data. The primary findings of our study are: (1) alternative and neural cholesterol clearance pathway of bile acid synthesis pathway genes were expressed consistently in the brain samples, indicating that these pathways are prevalent in the brain as compared to the classical bile acid synthesis pathway; (2) targeted metabolomics analysis of post-mortem brain samples identified primary and secondary bile acids and higher ratio of GCDCA:CA and secondary bile acids like DCA, LCA, TDCA, CDCA and GDCA in AD vs controls suggests that these bile acids might be associated with cognitive decline in AD; (3) the presence of secondary bile acids in metabolomics data suggests possible role of gut microbiome in AD and highlights the need to study the gut-brain axis to understand changes in AD; (4) transporters associated with taurine and cholesterol metabolism showed different usage based on our genome-scale metabolic network analysis of three independent cohorts; and (5) transcriptional regulatory network analysis identified transcription factors including *PPARA, RXRG* and *SREBF2* regulating bile acid and cholesterol genes in the brain.

### Role of bile acids in AD pathophysiology and use of genome-scale metabolic models

Bile acids are derived from cholesterol and their synthesis is regulated by complex feedback mechanisms^12,18^. Recent studies have identified bile acids in brain samples and linked them with cognitive decline in AD^7,10,19^. To understand the physiological role of bile acids in the brain of AD and CN individuals, we analyzed transcriptome data from post-mortem brain samples obtained from three independent cohorts and identified genes involved in the alternative bile acid pathway were expressed compared to the classical pathway in the brain. The alternative bile acid pathway is initiated by *CYP27A1* that catalyzes the steroid side-chain oxidation and in the subsequent step forms C24-bile acids. It is also known that cholesterol is converted to 24-hydroxycholesterol by sterol 24-hydroxylase (*CYP46A1*) in the brain and the gene was found to be expressed in the brain samples. The primary bile acids conjugate with glycine and taurine to form secondary bile acids. Taurine has a neuroprotective role in the brain and bile acids conjugated with taurine are found to be present in brain. Metabolomics data of serum samples showed AD patients had higher levels of serum taurine compared to controls, indicating taurine transport across the BBB might be affected in AD. The presence of secondary bile acids in the post-mortem brain samples suggests that these bile acids are either endogenously present in the brain or they are transported through the BBB. Bile acids such as ursodeoxycholic acid, taurocholic acid, taurolithocholic acid 3-sulfate and 3-dehydroxychenodeoxy cholic acid were also identified from our analysis and role of these bile acids can be probed further. Based on an association study, taurolithocholic acid was predicted to be a cytotoxic bile acid whereas chenodeoxycholic acid and ursodeoxycholic acid were predicted as neuroprotective bile acids^7^. Our analysis of transcriptome data of 2114 samples mapped into metabolic networks of brain regions implicated reactions involved in the production of metabolites such as 7α-hydroxycholesterol, 7α-hydroxy-5β-cholestan-3-one, 7α-hydroxycholestene-3-one and other derivatives that are formed through *CYP7A1* being significantly different (p-values for these reactions reported in Table 2) between AD vs CN. Although *CYP7A1* was not expressed in the post-mortem brain samples, the difference in abundance of these metabolites in AD vs CN suggests that we should explore the possibility of these metabolites entering the brain through the periphery. In this study, we have used transcriptomic data that was available from three independent cohorts. Transcriptomics data is insufficient to parametrize the metabolic models, but if a denser longitudinal omics data becomes available in the future it will help in improvising the predictions from these *in silico* models. Although, we identify reactions that are significant in these conditions, the directionality of the reactions can be solidly determined only if we have additional time-series metabolomics data (and ideally isotopic labeling experiments) to support these changes. Methods are now being developed to obtain cell type-specific data, so that we can gain additional information into the cells that are involved in regulating metabolic changes in AD. Generation of such data will help in refining the models and making more accurate predictions. Our brain-tissue metabolic models can be used by the community to capture *in silico* changes and possibly identify metabolic biomarkers prior to disease manifestation, making them useful in understanding interactions and mechanisms between different classes of metabolites and AD pathophysiology.

### Transcriptional regulation of bile acid and cholesterol genes

Metabolism is influenced by regulation of transcription factors and metabolic genes. In this study, we used a reconstruction transcriptional regulatory network of brain (and selected brain regions) to identify candidate TFs that may interact with genes in cholesterol and bile acid metabolism. We identified *SREBF2, PPARA, RXRG* and other transcription factors, some of which have been studied and implicated in Alzheimer’s disease. *SREBF2* expression enhances cholesterol levels^40^ and presumably oxysterol and cholestenoic acid levels which are ligands of LXR^41^. LXRs and the genes regulated by LXRs such as *ABCA1, ABCG1* and *APOE*, modulate intracellular cholesterol content and cholesterol efflux and have been associated with AD pathogenesis^42^. Our analysis also identified *PPARA* as putative regulator of *ABCA1* and recent studies have demonstrated that PPAR pathway activation increased *ABCA1* levels, that in turn lead to APOE lipidation and amyloid ß clearance^43^. We also identified transport of taurine as an important factor from the metabolic analysis. *SLC6A6* (neurotransmitter transporter, taurine) and *SLC36A1* (neutral amino acid/proton symporter) play a role in taurine transport. Although there was a 1.02 to 1.3-fold change in the expression of these transporters in AD compared with control samples of the three cohorts across four tested brain regions, this difference was only found to be statistically significant in cerebellum (Supplementaryxf file 1). Integration of expression data with metabolic network of brain regions identified reactions involving taurine transporters that were statistically significant in AD versus controls, further supporting their potential role. We had also *STAT1* is a putative TF of *SLC6A6*, identified from our analysis of brain regulatory network. Studies have suggested that the increased *STAT1* may be involved in inflammation in AD brain^37,44^. *NEUROD6* regulates the activity of *SLC36A1*, a proton-coupled amino acid transporter. *NEUROD6* is a basic helix-loop-helix TF and SNPs in *NEUROD6* have been associated with AD, especially in APOE4+ women^45^. Our analysis has been able to capture metabolic genes and putative TFs that regulate them. These findings can be further strengthened by generation of higher quality footprint data from brain samples.

### Studying the gut-brain axis to understand physiological changes in AD

Increasing evidence from experimental and clinical data suggests influence of gut-brain axis and gut microbiota in neurodegenerative diseases^46,47^. From our metabolic analysis we identified taurolithocholic, 3-dehydrochenodeoxycholic, and ursodeoxycholic acid, secondary bile acids, significant in AD compared to CN^48^ suggesting a possible connection to the gut microbiome. Recently, the bile acid deconjugation and biotransformation pathways have been reconstructed in a resource of genome-scale reconstructions of over 800 human gut microbes^49,50^. Of these, only 23 species could synthesize 3-dehydrochenodeoxycholic acid, only four could synthesize lithocholic acid, and only three could synthesize ursodeoxycholic acid^50^. For instance, the species *Ruminococcus* (*Blautia*) gnavus, and *Collinsella aerofaciens* synthesize 3-dehydroxychenodeoxycholic and ursodeoxycholic acid, and *Eggerthella lenta* synthesizes 3-dehydrochenodeoxycholic and several Clostridiales representatives synthesize lithocholic acid^50^, indicating these species may play a role in Alzheimer’s disease. Interestingly, increased lithocholic acid in plasma has recently been proposed as a potential biomarker for Alzheimer’s disease^51^. The personalized brain models developed in this study could be joined with personalized microbial community models established previously^50,52^. In future efforts, such combined host-microbe metabolic modeling will yield more insight into mechanisms underlying altered bile acid metabolism in Alzheimer’s disease.

## Methods

### 1. Transcriptome analysis of post-mortem brain samples

Transcriptome data was obtained from post-mortem brain samples of AD patients and cognitively normal individuals from Religious Orders Study and Memory and Aging Project (ROSMAP), Mayo Clinic, University of Florida, Institute for Systems Biology and Mount Sinai Brain Bank (MSBB). 265 samples of temporal cortex (TC) and cerebellum (CER), 632 samples of frontal cortex (FC), 303 samples of frontal pole (FP), superior temporal gyrus (STG), inferior frontal gyrus (IFG) and parahippocampal gyrus (PHG) with pathologies such as AD, MCI, Parkinson’s and control were analyzed and used for construction of brain region-specific metabolic models. ROSMAP data can be requested via the Rush Alzheimer’s Disease Center website (https://www.radc.rush.edu/). RNA-seq libraries were prepared by different methods such as poly-A enriched, strand-specific and ribo-zero. Table 1 has information of number of patients with various pathologies and controls and methods used for RNA-sequencing. The data used in the preparation of this article were downloaded from Synapse (https://www.synapse.org/#!Synapse:syn2580853/). We performed two-tailed t-test with Benjamini-Hochberg correction to identify differentially expressed genes with corresponding p-values. The differential expression analysis for transcriptome data from three independent cohort is presented in Supplementary file 1.

### 2. Bile acid sample preparation and analysis

Participants of the Religious Orders Study (ROS) are comprised of Catholic brothers, nuns, and priests who were cognitively normal at study entry and agreed to annual clinical examinations and brain donation at time of death. The Rush Memory and Aging Project (MAP) is a companion study that includes community-dwelling older adults that all agreed to evaluations similar to ROS. Quantification of bile acid concentration was performed at the University of Hawaii cancer center. The bile acid-free matrix (BAFM) was used to prepare bile acid calibrators. Extracts of brain tissue along with bile acid reference standards were subjected to instrumental analysis^53,54^. All of the 57 bile acid standards were obtained from Steraloids Inc. (Newport, RI) and TRC Chemicals (Toronto, ON, Canada) and 9 stable isotope-labeled standards were obtained from C/D/N Isotopes Inc. (Quebec, Canada) and Steraloids Inc. (Newport, RI). A Waters ACQUITY ultra performance LC system coupled with a Waters XEVO TQ-S mass spectrometer was used for all analyses. Chromatographic separations were performed with an ACQUITY BEH C18 column. UPLC-MS raw data obtained with negative mode were analyzed using TargetLynx™ applications manager to obtain calibration equations and the quantitative concentration (uM) of each bile acid. Bile acids were measured from the dorsolateral prefrontal cortex of 111 individuals with brain pathology (51 CN, 31 MCI and 27 AD at the time of death). Metabolomics data can be accessed with permission at https://www.synapse.org/#!Synapse:syn10235594. We calculated the ratio of primary and secondary bile acids measured in metabolomics study and performed two-tailed t-test to calculate p-value for each bile acid.

### 3. Brain region-specific metabolic reconstruction

We used transcriptome data (https://www.synapse.org/#!Synapse:syn2580853/) derived from post-mortem brain samples of three independent cohorts: Mayo clinic, ROSMAP and Mount Sinai Brain Bank. These cohorts contained information of different brain regions (CER, FC, TC, FP, STG, IFG and PHG) and the data was to generate brain region-specific metabolic networks. Transcriptome data was converted to binary by considering transcripts with values less than 25^th^ percentile in the matrix as 0 otherwise 1. We calculated ubiquity scores for genes in each brain region separately and used those for implementing mCADRE workflow^55^. The Recon 3D model^31^ of human metabolism was used as template to reconstruct brain region-specific metabolic networks as this model had information of reactions related to the primary and conjugated primary acids additionally added to refine the model. Once the draft reconstructions were generated, we used functions in COBRA toolbox to identify dead end metabolites and used reactions from Recon 3D model for removing gaps in the network. This step was carried out for each metabolic network reconstructed for brain regions. We also removed the reactions belonging to drug metabolism from the network, as they were not related to functions in the brain. Only partial urea cycle is reported to be active in the brain, and so we identified enzymes in the urea cycle that are present in brain^56^ and included the reactions related to these genes in the metabolic networks. The list of reactions is provided in Supplementary file 4-10. We did manual curation for genes present in the metabolic network using information from Human Protein Atlas^57^ for genes expressed in brain. This effort helped in providing further evidence for genes being present in the metabolic networks for brain regions. We also included metabolites defined in the cerebrospinal fluid (CSF) (by metabolomics data as well as the whole-body metabolism reconstruction^52^ and metabolites that can be taken up across the blood brain barrier (BBB) from blood into the CSF^58,59,60,61,62,63^. The list of metabolites that can pass BBB is provided in Supplementary file 19. We tested our models for 16 metabolic tasks (Supplementary file 11) that are brain specific and the models passed 50-70% of those tests, except for superior temporal gyrus metabolic network. As astrocytes are predominantly involved in maintaining brain physiology^64^, we used objective function of astrocytes for our brain metabolic networks. We constrained bounds of exchange reactions using information from a published work on metabolic interactions between cell types in brain^65^. Details of metabolites involved in objective function and bounds for constrained reactions are given in Supplementary file 4-10. We generated context-specific personalized metabolic networks for all samples included in our study using iMAT algorithm^66^ using brain region-specific reconstructions. Flux variability analysis^67^ was carried out to determine bounds for reactions in metabolic networks. We used COBRA toolbox v3.0^68^ for metabolic analysis that was implemented in MATLAB R2018a and academic licenses of Gurobi optimizer v7.5 and IBM CPLEX v12.7.1 were used to solve LP and MILP problems.

### 4. Reaction and pathway-level analysis

We carried out flux variability analysis^67^ for each context-specific personalized metabolic network and used the values for predicting metabolic changes in AD versus CN individuals and sex of the individuals. We created a matrix for reactions present in the personalized metabolic networks for each brain region and identified reactions that carried either minimum (*vmin*) or maximum (*vmax*) flux in the network. If a reaction carried flux it was assigned a state of 1, otherwise 0. This resulted in a matrix containing binary values for all reactions in 2114 context-specific personalized metabolic networks for seven brain regions. We used this scheme to classify the reactions and obtain information not only on the basis of flux measurements but also their activity in each network. Using Fisher’s exact test, we calculated p-values and those reactions with p-value < 0.05 were identified as significant reactions in these groups.

### 5. Metabolic regulatory network

The transcriptional regulatory network analysis (TReNA) package (https://rdrr.io/bioc/TReNA/) was used for identifying transcription factors (TFs) that are part of the co-expression modules of interest. Brain-specific transcriptional regulatory network was constructed^69^ using information from ENCODE. We downloaded the DNase Hypersensitivity (DHS) fastq files from ENCODE for all available brain samples and aligned the sequences using the SNAP method^70^. We performed two alignments using seed size of 16 and 20bp. The length of sequence data was >50 bp. The regions of open chromatin were identified using peak calling algorithm, F-Seq^71^. Footprints were generated using default parameters for Wellington^72^ and HINT^73^. Our method generated individual gene models and those footprints that are within the proximal promoter region (+/-5 kb of the transcription start site) are considered as priors in assessing the relationship between the expression of the TF and target genes. We prioritized putative TF regulators for each gene in the model using LASSO regression techniques, Pearson and Spearman correlation and random forest methods and projected the scores from these approaches into PCA space. The principal components were summed together to obtain a single composite score called pcaMax. This process is part of the trena package in Bioconductor (https://rdrr.io/bioc/TReNA/) and we applied the method to the post-mortem samples from temporal cortex from Mayo Clinic. We used metabolic genes identified from reaction-level analysis involved in bile acid and cholesterol metabolism and mapped top five transcription factors that interact with these metabolic genes and created an interaction network. This interaction networks gave information for transcription factors that regulate metabolic genes and are involved in significant reactions in AD versus cognitively normal individuals. Cytoscape 3.7.1^74^ was used for visualizing the brain transcriptional regulatory network.

## Data availability

Transcriptome data: https://www.synapse.org/#!Synapse:syn2580853/

Metabolomics data: https://www.synapse.org/#!Synapse:syn10235594

## Code availability

The code used for reconstruction and model generation and simulation are provided in GitHub (https://github.com/PriceLab/Bile_acid_AD)

## Funding

Funding for the ADMC (Alzheimer’s Disease Metabolomics Consortium, led by Rima Kaddurah-Daouk at Duke University) was provided by the National Institute on Aging (NIA) grant R01AG046171, a component of the Accelerated Medicines Partnership for AD (AMP-AD) Target Discovery and Preclinical Validation Project (https://www.nia.nih.gov/research/dn/amp-ad-target-discovery-and-preclinical-validation-project) and the National Institute on Aging grant RF1 AG0151550, a component of the M^2^OVE-AD Consortium (Molecular Mechanisms of the Vascular Etiology of AD– Consortium (https://www.nia.nih.gov/news/decoding-molecular-tiesbetween-vascular-disease-and-alzheimers). The Religious Orders and the Rush Memory and Aging studies were supported by the National Institute on Aging grants P30AG10161, R01AG15819, R01AG17917, and U01AG46152. Additionally, MA, RKD, and GK are supported by NIA grants RF1 AG058942 and R01 AG057452. MA and GK are also supported by funding from Qatar National Research Fund NPRP8-061-3-011. KN is supported by NIA grants NLM R01 LM012535 and NIA R03 AG054936. AJS is supported by NIH grants including P30 AG010133, R01 AG019771 and R01 CA129769. WJG is supported by funding from the UK Biotechnology and Biological Sciences Research Council (grant numbers BB/I001735/1 and BB/N015932/1) and the Engineering and Physical Sciences Research Council via an Impact Acceleration Account to Swansea University. JTY is supported by the Institute for Systems Biology’s Translational Research Fellows Program.

## Acknowledgements

The authors would like to thank the AMP-AD consortium for funding the project and AMP-AD Knowledge Portal for sharing the data. The authors would also like to thank members of the Hood-Price group at ISB for their support and help.

## Authors contribution

NDP and RKD conceived and led the study. PB reconstructed the brain region-specific metabolic networks; PB, CF, and JY analyzed the transcriptomics data for post-mortem brain samples downloaded from the AMP-AD knowledge portal; AKP analyzed the metabolomics data of brain; IT provided the list of metabolites that can cross blood-brain barrier; WJG provided valuable comments for bile acid analysis; WJ measured bile acids from post-mortem brain samples; AMP-AD consortium and the Alzheimer Disease Metabolomics Consortium collected transcriptomics and metabolomics data; PB, CF, JTY, JY, AKP, KN, AH, SM, GL, AJS, MA, GK, WJG, IT, RKD, and NDP contributed to the writing of this paper.

## Competing interest

The authors declare no competing interests.

## Notes

#### Summary of Updates

The revised version of the paper has updated figure numbers

